# Inhibition of ErbB kinase signalling promotes resolution of neutrophilic inflammation

**DOI:** 10.1101/738682

**Authors:** A. Rahman, K. M. Henry, K. D. Herman, A. A. R Thompson, H. M. Isles, C. Tulotta, D. Sammut, J. J. Y. Rougeot, N. Khoshaein, A. E. Reese, K. Higgins, C. Tabor, I. Sabroe, W. J. Zuercher, C. O. Savage, A. H. Meijer, M. K. B. Whyte, D. H. Dockrell, S. A. Renshaw, L. R. Prince

**Affiliations:** Department of Infection, Immunity and Cardiovascular Disease, University of Sheffield, Sheffield, UK; Department of Biochemistry and Molecular Biology, Faculty of Biological Sciences, University of Dhaka, Dhaka, Bangladesh; The Bateson Centre, University of Sheffield, UK; Institute of Biology, Leiden University, Leiden, The Netherlands; SGC-UNC, Division of Chemical Biology and Medicinal Chemistry, UNC Eshelman School of Pharmacy, University of North Carolina at Chapel Hill, USA; Immuno-Inflammation Therapy Area Unit, GlaxoSmithKline Research and Development Ltd., Stevenage, UK; MRC centre for Inflammation Research, University of Edinburgh, UK

**Author notes:** Joint corresponding authors: Lynne Prince and Stephen Renshaw. Currently at: Institute of Immunology and Immunotherapy, University of Birmingham, UK.

## Abstract

Neutrophilic inflammation with prolonged neutrophil survival is common to many inflammatory conditions, including chronic obstructive pulmonary disease (COPD). There are few specific therapies that reverse neutrophilic inflammation, but uncovering mechanisms regulating neutrophil survival is likely to identify novel therapeutic targets. Screening of 367 kinase inhibitors in human neutrophils and a zebrafish tail fin injury model identified ErbBs as common targets of compounds that accelerated inflammation resolution. The ErbB inhibitors gefitinib, CP-724714, erbstatin and tyrphostin AG825 significantly accelerated apoptosis of human neutrophils, including neutrophils from people with COPD. Neutrophil apoptosis was also increased in Tyrphostin AG825 treated-zebrafish *in vivo*. Tyrphostin AG825 decreased peritoneal inflammation in zymosan-treated mice, and increased lung neutrophil apoptosis and macrophage efferocytosis in a murine acute lung injury model. Tyrphostin AG825 and knockdown of *egfra* and *erbb2* by CRISPR/Cas9 reduced inflammation in zebrafish. Our work shows that inhibitors of ErbB kinases have therapeutic potential in neutrophilic inflammatory disease.

## Introduction

Neutrophilic inflammation is central to chronic inflammatory diseases such as rheumatoid arthritis and chronic obstructive pulmonary disease (COPD), which impose an increasing social and economic burden on our aging population. In these diseases, clearance of neutrophils by apoptosis is dysregulated, but to date it has not been possible to therapeutically modify this. The anti-inflammatory phosphodiesterase-4 inhibitor, roflumilast, targets systemic inflammation associated with COPD and reduces moderate to severe exacerbations in severe disease, possibly via effects on eosinophils (Martinez et al., 2018; Rabe et al., 2018). Recognising the urgent need for new therapies, we interrogated neutrophil inflammation and survival pathways using an unbiased approach focusing on potentially druggable kinases. Neutrophil persistence in tissues, caused by a delay in apoptosis, can result in a destructive cellular phenotype, whereby neutrophils have greater potential to expel histotoxic factors such as proteases and oxidative molecules onto surrounding tissue. This can occur either actively (by degranulation) or passively (by secondary necrosis). In COPD, among other diseases, delayed apoptosis is considered to be a key part of the pathogenesis, occurring either as a result of pro-survival factors that are present in the lung microenvironment or an innate apoptosis defect (Brown, Elborn, Bradley, & Ennis, 2009; Haslett, 1999; Pletz, Ioanas, de Roux, Burkhardt, & Lode, 2004; J. Zhang, He, Xia, Chen, & Chen, 2012). Despite this mechanistic understanding, there are no effective treatment strategies in clinical use to specifically reverse this cellular mechanism.

Accelerating neutrophil apoptosis has been shown to promote the resolution of inflammation in multiple experimental models (Burgon et al., 2014; Chello et al., 2007; Ren et al., 2008; Rossi et al., 2006). A number of studies highlight the importance of protein kinases in regulating neutrophil apoptosis (Burgon et al., 2014; Rossi et al., 2006; Webb et al., 2000) and therefore reveal potential therapeutically targetable pathways for inflammatory disease. A growing class of clinically-exploited small molecule kinase inhibitors are being intensively developed (Wu, Nielsen, & Clausen, 2015), making this a timely investigation. Using parallel unbiased screening approaches *in vitro* and *in vivo,* we here identify inhibitors of the ErbB family of receptor tyrosine kinases (RTKs) as potential therapeutic drivers of inflammation resolution. The ErbB family consist of four RTKs with structural homology to the human epidermal growth factor receptor (EGFR/ErbB1/Her-1). In an *in vivo* zebrafish model of inflammation, we show that inhibition of ErbBs, pharmacologically and genetically, reduced the number of neutrophils at the site of injury. Furthermore, ErbB inhibitors reduced inflammation in a murine peritonitis model and promoted neutrophil apoptosis and clearance by macrophages in the mouse lung. This study reveals an opportunity for the use of ErbB inhibitors as a treatment for chronic neutrophilic inflammatory disease.

## Results

### Identifying kinases regulating the resolution of neutrophilic inflammation *in vivo*

Using a well-characterised transgenic zebrafish inflammation model (Henry, Loynes, Whyte, & Renshaw, 2013; Renshaw et al., 2006), we adopted a chemical genetics approach, which has great potential for accelerated drug discovery (Jones & Bunnage, 2017). We initiated inflammation by controlled tissue injury of the zebrafish tail fin and screened a library of kinase inhibitors in order to establish which kinases could be exploited to enhance inflammation resolution *in vivo* (Fig. S1A). We quantified the ability of a library of 367 publicly available kinase inhibitors (PKIS) (Elkins et al., 2016) to reduce neutrophil number at the site of injury during the resolution phase of inflammation. The screen identified 16 hit compounds which reduced neutrophil number at the site of injury in the zebrafish model (Fig. 1A). For each compound the degree of kinase inhibition had been established (Elkins et al., 2016) (Fig. 1A). A number of kinases were inhibited by the 16 compounds, with Abelson murine leukaemia viral homolog 1 (ABL1), Platelet-derived growth factor receptor (PDGFR) α, PDGFRβ, p38α and ErbB4 being the top five most frequently targeted kinases overall. In addition to frequency of target, we also interrogated selectivity of compound. The most selective compounds, i.e. those that strongly inhibited individual kinases or kinase families, targeted the kinases YES, ABL1, p38 and the ErbB family. Apoptosis is an important mechanism contributing to inflammation resolution; we therefore sought to identify kinases common to both inflammation resolution and neutrophil apoptosis pathways.

**Figure 1:**
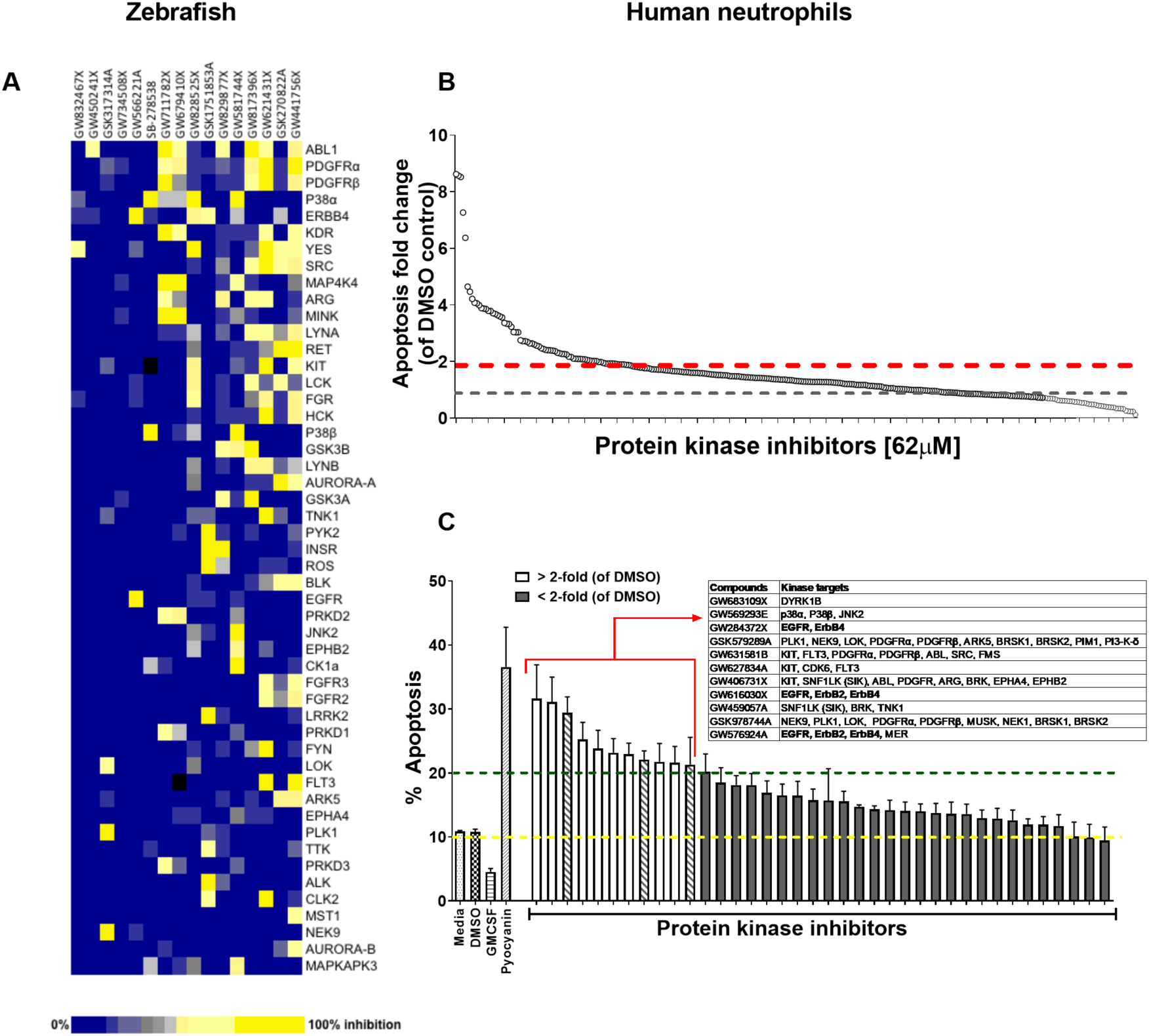
A protein kinase inhibitor compound library screen identifies compounds that promote the resolution of inflammation *in vivo* and neutrophil apoptosis *in vitro*. (A) *mpx:GFP* zebrafish larvae (3 dpf) that had undergone tail fin transection resulting in an inflammatory response at 6 hpi were incubated with individual PKIS compounds [25µM] 3 larvae/well for a further 6h. Wells were imaged and manually scored between 0-3 on the basis of GFP at the injury site in the larvae. ‘Hit’ compounds scored ≥ 1.5 (n=2, 3 larvae per compound per experiment). Publicly available kinase profiling information was generated previously by Elkins *et al*. (2016) and kinase inhibition of each compound [1µM] is shown as a gradient of blue to yellow. Hit compounds were ranked horizontally (left to right) from the most to least selective. Kinases (listed on the right) were vertically ranked from top to bottom from the most to least commonly targeted by inhibitors in PKIS. (B) PKIS compounds were incubated with primary human neutrophils for 6h. The entire library, at [62µM], was screened on 5 separate days using 5 individual donors. Apoptosis was assessed by Annexin-V/TO-PRO-3 staining by flow cytometry and the percentage apoptosis calculated as Annexin-V single plus Annexin-V/TO-PRO-3 dual positive events. Data are expressed as fold change over DMSO control and each circle represents a single compound. Sixty two compounds accelerated apoptosis ≥ 2 fold as identified by red dotted line (n=1). Grey dotted line represents level of apoptosis in DMSO control (i.e. no change). (C) Of the 62 compounds identified above, 38 of the most specific inhibitors were incubated with neutrophils at [10µM] for 6h and apoptosis measured as above. Controls included media, DMSO, GMCSF [50 u/mL] and pyocyanin [50µM]. Eleven compounds (white bars) accelerated apoptosis ≥ 2 fold over DMSO control (as identified by dotted line). Kinases targeted by the 11 compounds are shown in the inset table. Hatched bars represent data points in which ErbB inhibitors were used. Data are expressed as percentage apoptosis ± SEM, n=3 neutrophil donors.

### Identifying kinases regulating neutrophil apoptosis *in vitro*

Circulating neutrophils have a short half-life *in vivo* (Summers et al., 2010) and undergo spontaneous apoptosis in the absence of growth factors *in vitro*. We re-screened PKIS library compounds in a human neutrophil apoptosis assay for their ability to accelerate apoptosis (Fig. S1B). PKIS compounds were screened at 62µM in order to maximise the chance of identifying ‘hits’ and resulted in 62 compounds that accelerated neutrophil apoptosis ≥2-fold compared to DMSO control (Fig. 1B and Table S1). Secondary screening of top 38 compounds (chosen from the 62 hits based on greatest selectivity for kinase targets) was carried out at 10µM in order to reduce false positives. This yielded 11 compounds that accelerated neutrophil apoptosis ≥2-fold over control (as indicated by dashed green line, Fig. 1C). Kinases targeted by these compounds included DYRK1B, KIT, EGFR, ErbB2 & ErbB4, PDGFR, CDK6 and p38 (Fig. 1C, inset). The identification of known regulators of neutrophil survival (p38, PI3K) was encouraging support for the screen design and execution. We found that members of the ErbB family of RTKs were the next most frequently inhibited kinase family, being targeted by 3 highly selective compounds out of the 11 hits (Fig. 1C, inset). Since inhibitors of the ErbB family were common hits in both zebrafish and human screens, we hypothesised that targeting ErbBs may be a potential strategy to reduce inflammation.

### ErbB inhibitors accelerate neutrophil apoptosis

To address a role for ErbB antagonists in regulating neutrophil apoptosis we tested a range of clinical and non-clinical ErbB-targeting compounds. We show that among inhibitors of ErbBs that are in clinical use, the EGFR inhibitor, gefitinib, is the most effective in promoting neutrophil apoptosis, reaching significance at 50µM (Fig. 2A). The ErbB2-selective inhibitor, CP-724714 (Jani et al., 2007) also promoted neutrophil apoptosis in a dose-dependent manner (Fig. 2B) as did Erbstatin and tyrphostin AG825, selective for EGFR and ErbB2 respectively (Osherov, Gazit, Gilon, & Levitzki, 1993; Umezawa & Imoto, 1991) (Fig. 2C-D). Since caspase-dependent apoptosis is an anti-inflammatory and pro-resolution form of cell death, engagement of the apoptosis programme was verified biochemically by measuring phosphatidylserine (PS) exposure by Annexin-V staining (Fig. S2A-C). Furthermore, the pan-caspase inhibitor Q-VD-OPh (Wardle et al., 2011) completely abrogated Erbstatin and tyrphostin AG825-driven neutrophil apoptosis, confirming the caspase dependence of inhibitor mediated cell death (Fig. S2D-E).

**Figure 2:**
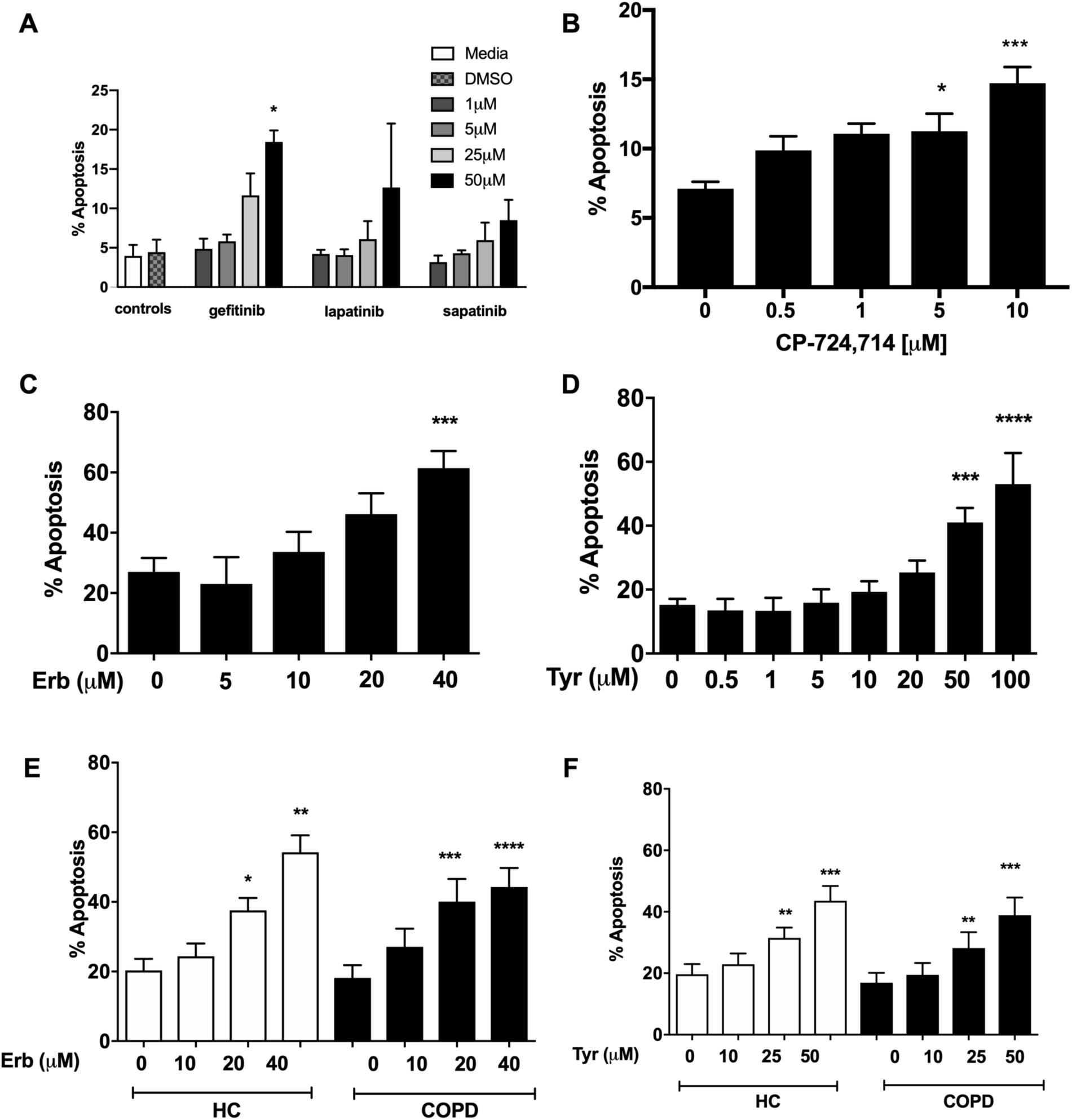
Inhibition of EGFR and ErbB2 drives apoptosis of neutrophils isolated from COPD patients and healthy subjects. Neutrophils were incubated with media or a concentration range of gefitinib (A), lapatinib (A), sapatinib (A), CP-724714 (B), erbstatin (Erb, C) or tyrphostin AG825 (Tyr, D) for 6h. Stars represent significant difference compared to DMSO control (indicated by “0” in B-D). Neutrophils from COPD patients (open bars) and age-matched healthy control subjects (black bars) were incubated with DMSO or a concentration range of erbstatin (E) or tyrphostin AG825 (F) for 6h. Apoptosis was assessed by light microscopy. The data are expressed as mean percentage apoptosis ± SEM from 3 (B, D), 4 (A,C), 10 (E,F COPD), or 7 (E,F HC) independent experiments using different neutrophil donors. Statistical significances between control and inhibitor was calculated by two-way ANOVA (A) or one-way ANOVA (B-F) with appropriate post-test, indicated as *p<0.05, **p<0.01, ***p<0.001, ****p<0.0001.

COPD is a chronic inflammatory disease associated with functionally defective circulating neutrophils, including a resistance to undergoing apoptosis during exacerbations (Pletz et al., 2004; Sapey et al., 2011). To show ErbB inhibition is effective in driving apoptosis in subjects with systemic inflammation, we isolated neutrophils from the blood of patients with COPD and age-matched healthy control subjects. Erbstatin and tyrphostin AG825 significantly increased apoptosis of neutrophils from both COPD patients and healthy control subjects in a dose dependent manner at both 6h (Fig. 2E-F) and 20h (data not shown).

ErbB inhibition overcomes neutrophil survival stimuli. Neutrophils are exposed to multiple pro-survival stimuli at sites of inflammation, which could undermine the therapeutic potential of anti-inflammatory drugs. Factors that raise intracellular cAMP concentration ([cAMP]_i_) are present during inflammation, and elevated [cAMP]_i_ is known to prolong neutrophil survival via activation of cAMP-dependent protein kinases (Krakstad, Christensen, & Doskeland, 2004; Vaughan et al., 2007). We show that erbstatin and tyrphostin AG825 significantly reversed N^6^-monobutyryl-cAMP (N^6^-MB-cAMP)-mediated survival (Fig. 3A-B). Similar effects were observed in neutrophils from patients with COPD (Fig. 3C). GMCSF is a key neutrophil chemoattractant and pro-survival factor, and is closely associated with the severity of inflammation in disease (Klein et al., 2000; Wicks & Roberts, 2016). We show that erbstatin and tyrphostin AG825 prevent GMCSF-mediated survival in COPD and age-matched healthy control neutrophils (Fig. 3D-E). GMCSF is known to promote neutrophil survival via the phosphatidylinositol 3-kinase (PI3K)/AKT pathway, ultimately leading to the stabilisation of the anti-apoptotic Bcl-2 family member, Mcl-1 (Derouet, Thomas, Cross, Moots, & Edwards, 2004; Klein et al., 2000). To investigate potential mechanisms underpinning the ability of tyrphostin AG825 to prevent GMCSF-mediated survival, we assessed AKT-phosphorylation as a measure of PI3K activation and found that tyrphostin AG825 reduced GMCSF-induced AKT phosphorylation after 15 and 30 min of treatment (Fig. 3F). Tyrphostin AG825 accelerated the spontaneous downregulation of Mcl-1 and also prevented GMCSF-induced stabilisation of Mcl-1 (Fig. 3G). These data show ErbB inhibition engages neutrophil apoptosis even in the presence of inflammatory stimuli and therefore has the potential to drive apoptosis at inflammatory sites.

**Figure 3:**
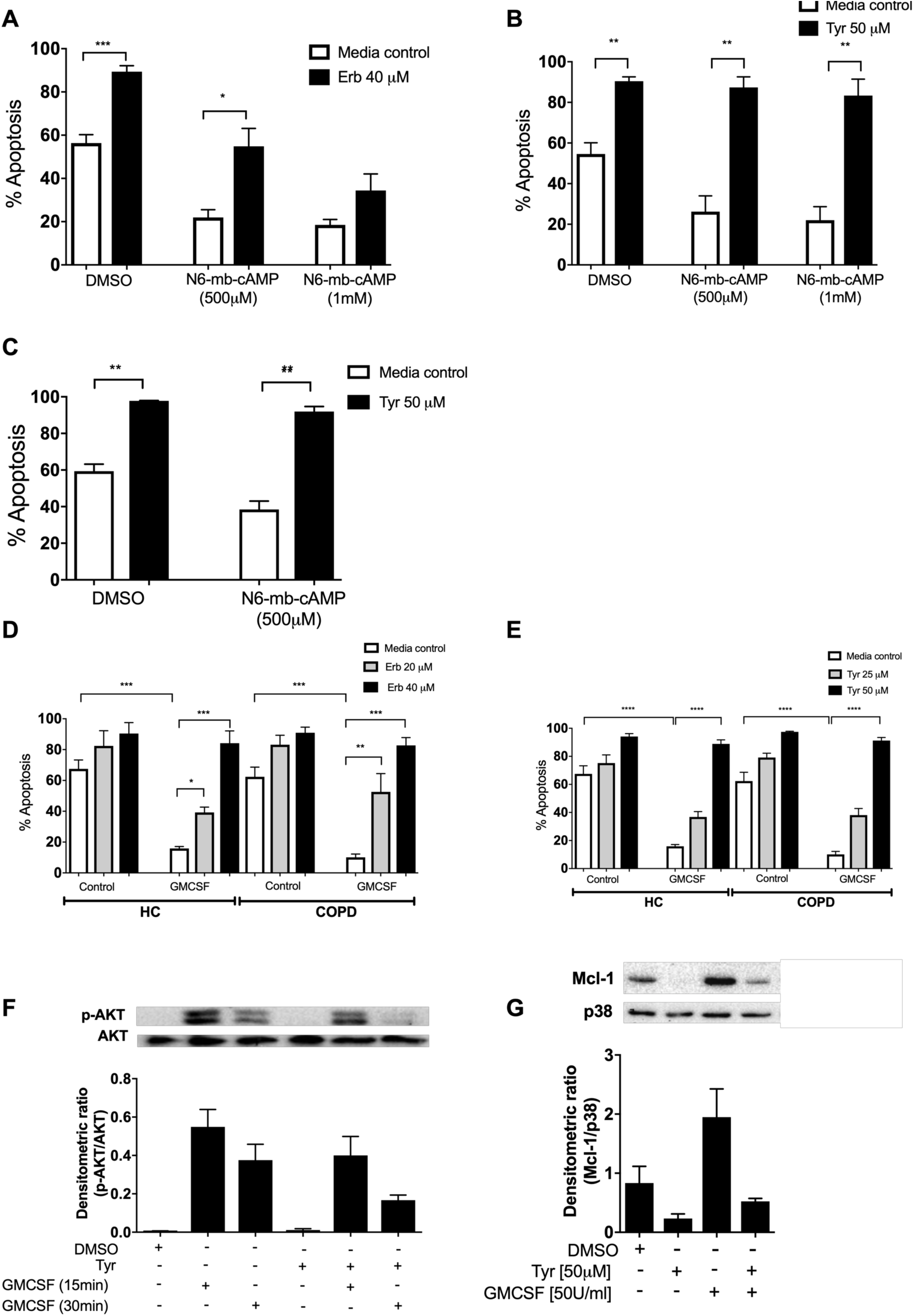
Erbstatin and tyrphostin AG825 overcome pro-survival effects of N^6^-MB-cAMP and GMCSF. Neutrophils were incubated with DMSO, Erbstatin [Erb, 40 µM] (A) or tyrphostin AG825 [Tyr, 50µM] (B) in the presence of DMSO or N^6^-MB-cAMP [500µM and 1mM] for 20h. Neutrophils isolated from COPD patients were incubated with DMSO or tyrphostin AG825 [50µM] in the presence of DMSO or N^6^-MB-cAMP [500µM] for 20h (C). Neutrophils isolated from COPD patients and age-matched healthy control subjects (HC) were incubated with DMSO, erbstatin (D) [20, 40µM] or tyrphostin AG825 (E) [25, 50µM] in the presence or absence of GMCSF [50u/mL] for 20h. Apoptosis was assessed by light microscopy. The data are expressed as mean percentage apoptosis ± SEM from 4-6 independent experiments. Statistical significances were calculated by one-way ANOVA with appropriate post-test and indicated as *p<0.05, **p<0.01, ***p<0.001. (F) Neutrophils were incubated with DMSO or tyrphostin AG825 [Tyr, 50µM] for 60 min before the addition of GMCSF [50u/mL] for 15 or 30 mins. (G) Neutrophils were incubated with DMSO, tyrphostin AG825 [50µM] for 60 min before the addition of GMCSF [50u/mL] for a further 7h. Cells were lysed, subjected to SDS-PAGE electrophoresis and membranes probed for p-AKT, Mcl-1 or loading controls, AKT and P38. Images are representative of 3 independent experiments. Charts show densitometric values of 3 individual immunoblots and are expressed as a ratio of target (p-AKT or Mcl-1) over loading control (AKT or P38, respectively).

Kinase microarray profiling reveals ErbB2 is phosphorylated by neutrophil survival stimuli. To explore whether ErbB family members are phosphorylated in response to survival stimuli we studied the activated kinome in human neutrophils stimulated with N^6^-MB-cAMP (Vaughan et al., 2007). A Kinex™ antibody microarray was performed to detect the phosphorylation of over 400 kinases and kinase-associated proteins and this data set was interrogated to seek evidence of activation of ErbB by N^6^-MB-cAMP. Of the phospho-specific antibodies, 17 yielded an increase over baseline control of ≥ 1.5 at 30 min and 8 at 60 min (Table 1). Among these targets, ErbB2 phosphorylation was detected at 30 min (1.94 > control) and 60 min (1.53 > control, Table 1). This suggests that ErbB is part of the neutrophil signalling response to survival stimuli. In support of this, we detected the presence of ErbB2 mRNA in human neutrophils by RT-PCR (Fig. S3) and a 60kD protein (Guillaudeau et al., 2012; Siegel, Ryan, Cardiff, & Muller, 1999; T. M. Ward et al., 2013), which was upregulated by GMCSF and dbcAMP (Fig. S3). ErbB3 was also detected in human neutrophils by ELISA (Fig. S3), at levels similar to those observed in other tissues in literature (Buta et al., 2016). We found ErbB3 expression was not regulated by growth factors, which may in part be due to regulation being primarily at the post-translational level.

**Table 1:**
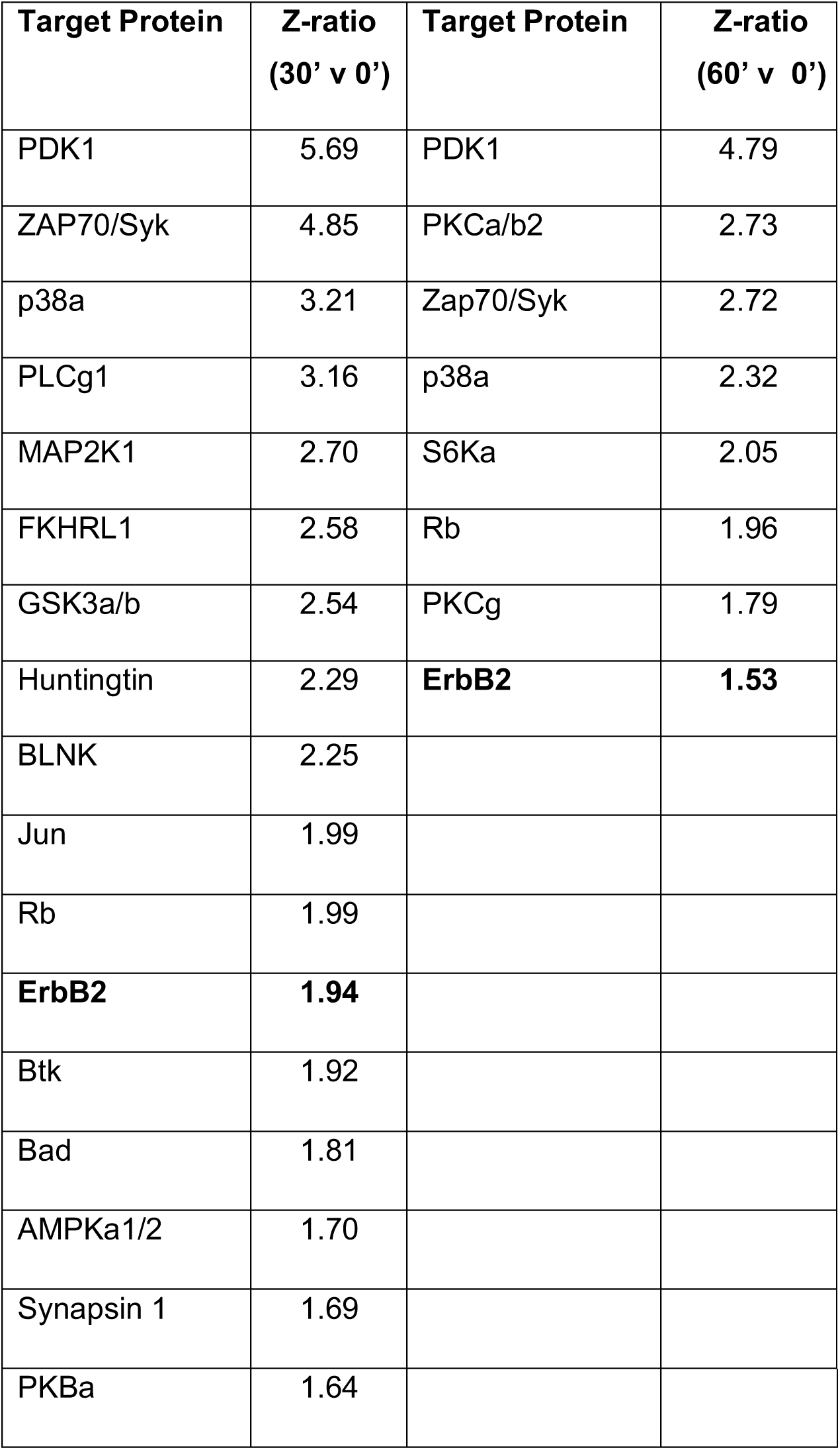
Kinexus antibody microarray analysis. Ultrapurified neutrophils were incubated with N^6^-MB-cAMP [100µM] for 30 and 60 min or lysed immediately following isolation (0’). Lysates from four donors were pooled prior to Kinex antibody microarray analysis. Table shows all targets for which phospho-antibodies had Z ratios of >1.5 compared to t=0 baseline control, at each timepoint. ErbB related antibodies are in bold.

| Target Protein | Z-ratio<br>(30' v 0') | Target Protein | Z-ratio<br>(60' v 0') |
| --- | --- | --- | --- |
| PDK1 | 5.69 | PDK1 | 4.79 |
| ZAP70/Syk | 4.85 | PKCa/b2 | 2.73 |
| p38a | 3.21 | Zap70/Syk | 2.72 |
| PLCg1 | 3.16 | p38a | 2.32 |
| MAP2K1 | 2.70 | S6Ka | 2.05 |
| FKHRL1 | 2.58 | Rb | 1.96 |
| GSK3a/b | 2.54 | PKCg | 1.79 |
| Huntingtin | 2.29 | **ErbB2** | **1.53** |
| BLNK | 2.25 |  |  |
| Jun | 1.99 |  |  |
| Rb | 1.99 |  |  |
| **ErbB2** | **1.94** |  |  |
| Btk | 1.92 |  |  |
| Bad | 1.81 |  |  |
| AMPKa1/2 | 1.70 |  |  |
| Synapsin 1 | 1.69 |  |  |
| PKBa | 1.64 |  |  |

### ErbB inhibitors and genetic knockdown increase apoptosis and reduce neutrophil number at the site of inflammation *in vivo*

To determine the ability of ErbB inhibition to exert an effect on neutrophil number and apoptosis *in vivo,* we used three complementary animal models of acute inflammation. To specifically address whether tyrphostin AG825 was able to accelerate apoptosis of neutrophils in the mammalian lung, we used a murine model of LPS-induced airway inflammation (Thompson et al., 2014). C57BL/6 mice nebulised with LPS developed an acute pulmonary neutrophilia after 48h, to a degree seen previously (Fig. 4A-B) (Thompson et al., 2014). Tyrphostin AG825 had no effect on percentage of, or absolute number of neutrophils or macrophages compared to DMSO control (Fig. 4A-B). Tyrphostin AG825 significantly increased the percentage of neutrophil apoptosis, both visualised as ‘free’ apoptotic cells (closed circles) and as a summation of both free apoptotic cells and apoptotic inclusions within macrophages in order to capture those that had been efferocytosed (closed triangles, Fig. 4C). Macrophage efferocytosis was also significantly elevated by tyrphostin AG825, compared to vehicle control (Fig. 4D), determined by counting the number of macrophages containing apoptotic inclusions as a proportion of total macrophages (Fig. 4E). We next tested the anti-inflammatory potential of tyrphostin AG825 when administered once inflammation was established, which is more representative of the clinical scenario. Mice were i.p injected with zymosan to induce peritonitis and after 4h were treated (i.p.) with tyrphostin AG825 or vehicle control. Total cell counts in peritoneal lavage were 2.2 x10^6^ in PBS vs 1.7 x10^7^ in zymosan treated animals at 4h demonstrating established inflammation at this time point (Navarro-Xavier et al., 2010). Importantly, tyrphostin AG825 does not induce leukopenia (Fig. 4F), however significantly fewer inflammatory cells were found in peritoneal lavage following tyrphostin AG825 treatment (Fig. 4G). The neutrophil chemoattractant and proinflammatory cytokine, KC, was reduced in tyrphostin AG825 treated mice, and concomitant with this, a trend for less IL-6 was also observed (Fig. 4H). IgM, which correlates with the number and activation of peritoneal B lymphocytes (Almeida et al., 2001), is significantly reduced in tyrphostin AG825-treated mice (Fig. 4I).

**Figure 4:**
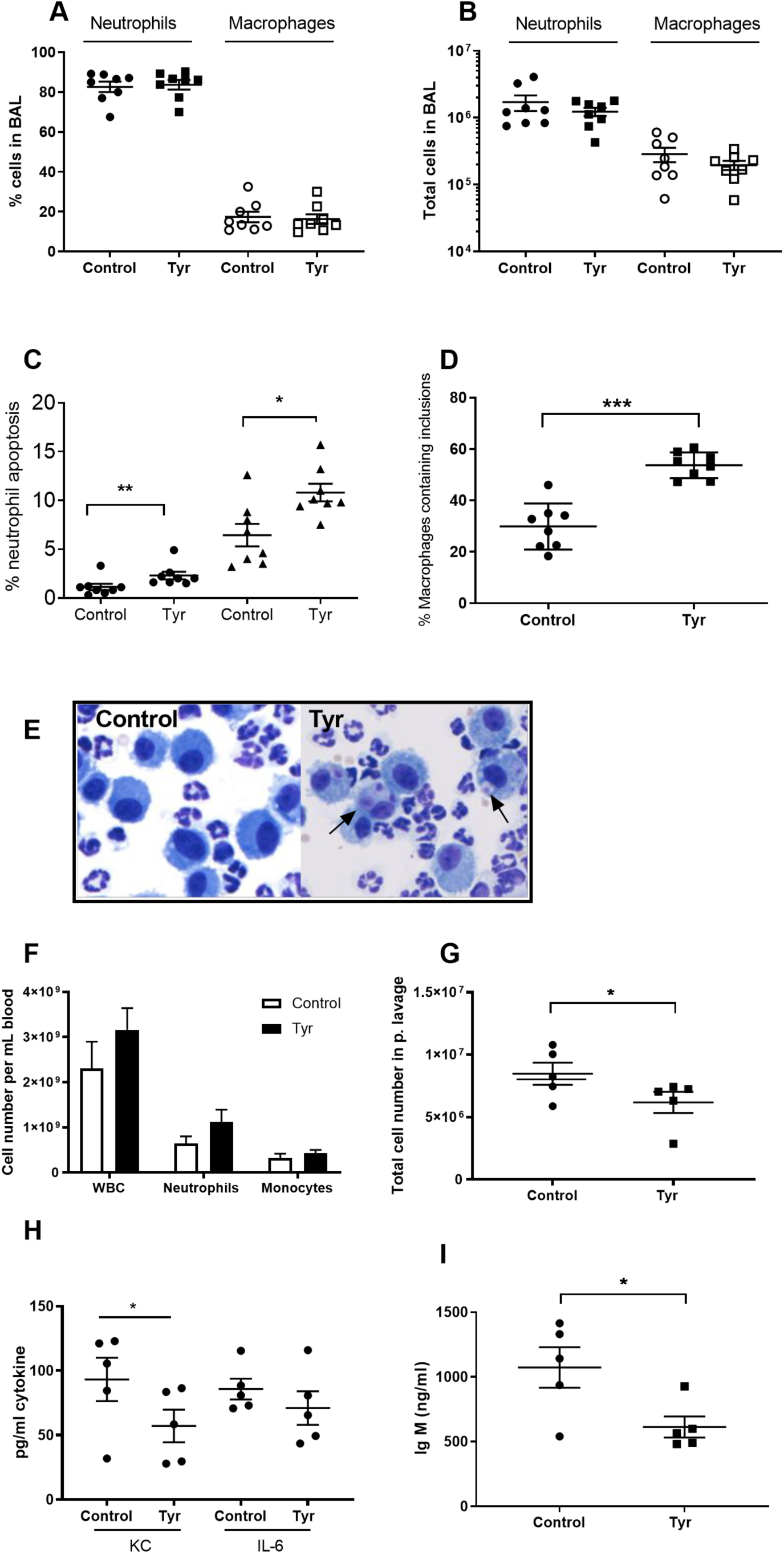
Tyrphostin AG825 increases neutrophil apoptosis and reduces inflammation in murine models of inflammation. C57BL/6 mice were nebulized with LPS and immediately injected intraperitoneally with either 10% DMSO (control, n=8) or 20mg/Kg tyrphostin AG825 (Tyr, n=8). After 48h the mice were sacrificed and subjected to bronchoalveolar lavage. Percentage neutrophils (A, closed icons) and macrophages (A, open icons) and absolute numbers of neutrophils (B, closed icons) and macrophages (B, open icons) in BAL were calculated by haemocytometer and light microscopy. (C) Percentage neutrophil apoptosis (circles) and percentage neutrophil apoptosis calculated by also including numbers of apoptotic inclusions visualised within macrophages (triangles) was assessed by light microscopy. (D) Macrophages containing 1 or more apoptotic inclusions expressed as a percentage of all macrophages. Light microscopy image showing apoptotic inclusions within macrophages as indicated by black arrows (E). C57BL/6 mice were injected i.p. with 1 mg zymosan and 4 h later injected i.p. with 20mg/Kg tyrphostin AG825 (Tyr, n=5) or 10% DMSO (Control, n=5). At 20h mice were sacrificed and subjected to peritoneal lavage. (F) WBC, neutrophils and macrophages in blood were measured by a Sysmex cell counter. Total cells in peritoneal lavage were counted by flow cytometry (G) and KC, IL-6 (H) and IgM (I) measured in lavage by ELISA. At least 2 independent experimental replicates each processing 1-3 mice/group were performed. Statistical significance was calculated by non-parametric t-test (Mann–Whitney U test), *p<0.05, **p<0.01, ***p<0.001.

To further extend this observation, we tested the ability of ErbB inhibitors to modulate neutrophilic inflammation resolution as a whole, in a model which encompasses multiple mechanisms of neutrophil removal including both apoptosis and reverse migration. In the *mpx*:GFP zebrafish tail fin injury model (Renshaw et al., 2006) (Fig. 5A) we were able to show that tyrphostin AG825 (Fig. 5B) and CP-724714 (Fig. 5C) significantly reduced the number of neutrophils at the site of injury at 4 and 8 hpi. Simultaneous gene knockdown of *egfra* and *erbb2* via CRISPR/Cas9 (referred to as ‘crispants’) also recapitulated this phenotype (Fig. 5D). Tyrphostin AG825 did not affect total neutrophil number (Fig. 5E), but *egfra* and *erbb2* crispants had significantly fewer neutrophils (Fig. 5F). As demonstrated by TSA and TUNEL double staining (Fig. 5G), tyrphostin AG825 upregulated neutrophil apoptosis at both the site of injury (Fig. 5H) and in the caudal hematopoietic tissue (CHT) of zebrafish (Fig. 5I). CHT neutrophil counts were unchanged between conditions (data not shown). *egfra* and *erbb2* crispants had increased numbers of apoptotic neutrophils at the site of injury, but this was not significant (Fig. 5J), perhaps suggesting the presence of compensatory mechanisms. These findings show that inhibiting ErbB RTKs accelerate neutrophil apoptosis *in vitro* and *in vivo* and enhance inflammation resolution, making ErbB inhibitors an attractive therapeutic strategy for inflammatory disease.

**Figure 5:**
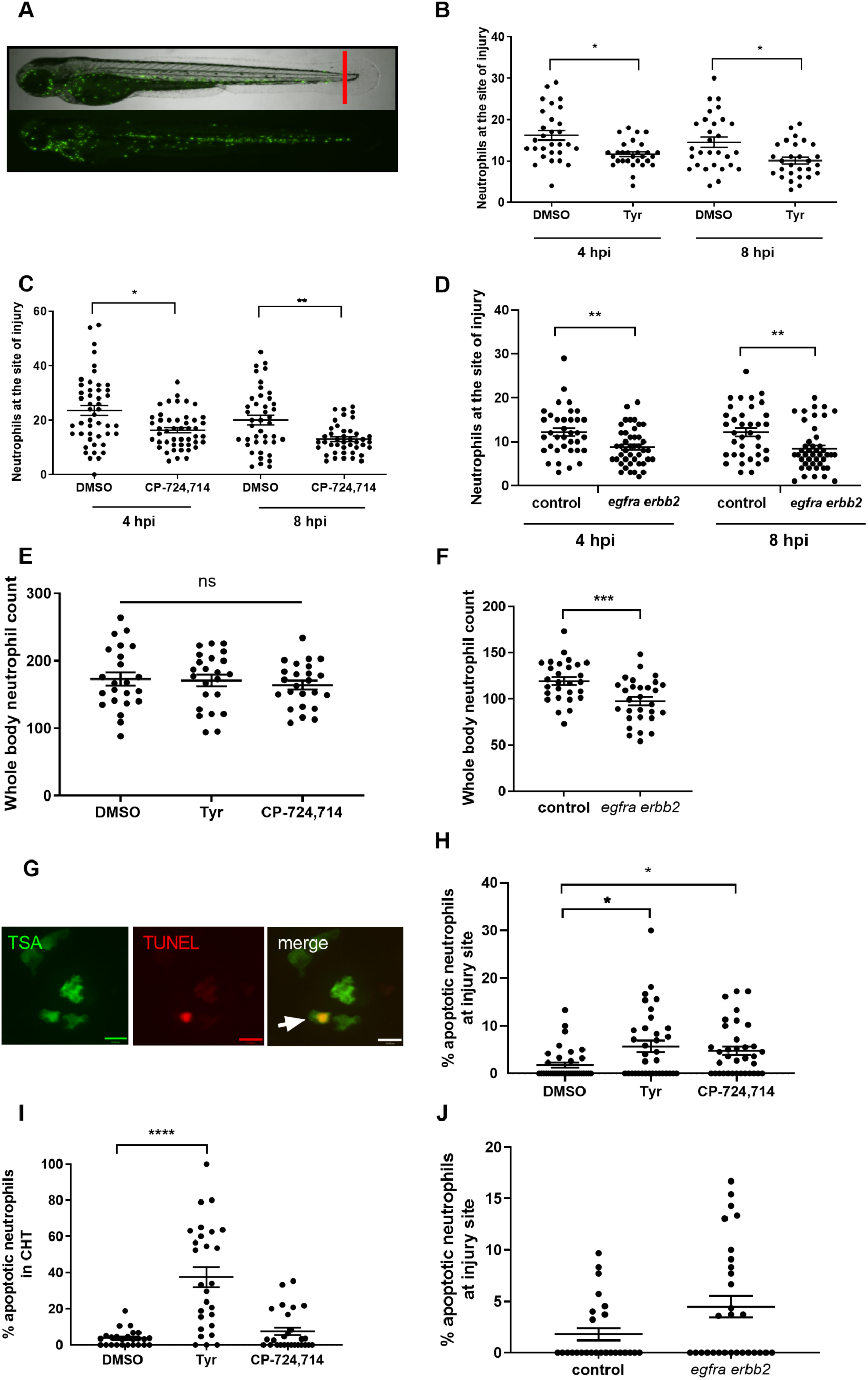
Pharmacological inhibition and genetic knockdown of *egfra* and *erbb2* by CRISPR/Cas9 reduces neutrophil number at the site of injury in a zebrafish model of inflammation. Tail fin transection was performed as indicated by the red line (A, upper image). Zebrafish larvae (*mpx*:GFP) were pre-treated at 2 dpf with DMSO, tyrphostin AG825 [Tyr, 10µM] (B, minimum n=28 larvae per condition), or CP-724714 [10µM] (C, minimum n=42 larvae per condition) for 16h followed by injury. *egfra* and *erbb2* crispants were generated and injured at 2 dpf (D, minimum n=36 larvae per condition). The number of neutrophils at the site of injury was determined at 4 and 8 hpi by counting GFP-positive neutrophils. To enumerate neutrophils across the whole body, uninjured inhibitor treated larvae (3 dpf) (E, minimum n=23 larvae per condition) or crispants (2 dpf) (F, minimum n=28 larvae per condition) were imaged by fluorescent microscopy (A, lower image). Apoptosis was measured at the site of injury after 8 hours by TSA and TUNEL double staining (G) (white arrow indicates TUNEL positive neutrophil, scale bar 10μM) of mpx:*GFP* tyrphostin AG825 [Tyr, 10µM] or CP-724714 [10µM] treated larvae at 3 dpf (H, minimum n=35 larvae per condition). Uninjured inhibitor treated larvae were assessed for neutrophil apoptosis in the CHT at 3 dpf (I, minimum n=27 larvae per group). Apoptosis at the tail fin injury site of *egfra erbb2* crispants at 2 dpf was also measured at 8 hpi (J, minimum n=26 larvae per group). All data collated from at least 3 independent experiments, displayed as mean ± SEM. Each icon shows one data point from one individual larvae. Statistical significances were calculated by two-way ANOVA (B-D) or one-way ANOVA (E, H, I) with appropriate post-test, unpaired-t test (F), Kruskal-Wallis test (I) with appropriate post-test or Mann-Whitney U test (J), and indicated as *p<0.05, **p<0.01, ***p<0.001, ****p<0.0001.

## Discussion

Neutrophils are powerful immune cells because of their destructive anti-microbial contents. A deleterious by-product of this is their remarkable histotoxic potential to host tissue, ordinarily held in check by the onset of apoptosis. The inappropriate suppression of neutrophil apoptosis underpins a number of chronic inflammatory diseases, and we are yet to have available an effective treatment strategy that can reverse this cellular defect in clinical practice. Here we show in human, mouse and zebrafish models of inflammation and neutrophil cell death that targeting the ErbB family of RTKs regulates neutrophil survival and resolves inflammation.

Promoting neutrophil apoptosis is a desirable approach for the resolution of inflammation, since apoptosis functionally downregulates the cell, promotes rapid cell clearance by efferocytosis and engages an anti-inflammatory phenotype in phagocytosing cells (Savill et al., 1989; Whyte, Renshaw, Lawson, & Bingle, 1999). As proof of principle, driving apoptosis experimentally promotes the resolution of inflammation across multiple disease models (Chello et al., 2007; Ren et al., 2008; Rossi et al., 2006). Several compounds targeting the ErbB family have been approved as medicines for the treatment of cancer (Singh, Attri, Gill, & Bariwal, 2016). Our findings open up the possibility of repurposing well-tolerated ErbB inhibitors for patients with inflammatory disease, potentially addressing a currently unmet clinical need.

The ErbB family are critical regulators of cell proliferation and are associated with the development of many human malignancies (Roskoski, 2014). In addition to the development of cancer, ErbB members have known roles in inflammatory diseases of the airway, skin and gut (Davies, Polosa, Puddicombe, Richter, & Holgate, 1999; Finigan et al., 2011; Frey & Brent Polk, 2014; Hamilton et al., 2003; Pastore, Mascia, Mariani, & Girolomoni, 2008). In the context of lung inflammation, ErbB2 is upregulated in whole lung lysates in murine bleomycin models of lung injury and EGFR ligands are increased in BAL from acute lung injury patients receiving mechanical ventilation (Finigan et al., 2011), suggesting ErbB signalling axes may play a role in the process of airway inflammation *in vivo*. We show, in murine models where Tyrphostin AG825 was administered either at the time of inflammatory stimulus or once inflammation was established, an impact on cell number, proinflammatory cytokine production and neutrophil apoptosis, further validating the use of ErbB inhibitors to reduce inflammation. The benefit of EGFR inhibitors in reducing inflammation in ventilator-induced and OVA/LPS-induced lung injury rodent models is shown by others, further supporting the targetting of this pathway in inflammatory disease settings (Bierman, Yerrapureddy, Reddy, Hassoun, & Reddy, 2008; Shimizu et al., 2018; Takezawa, Ogawa, Shimizu, & Shimizu, 2016).

Others have reported that neutrophils express members of the ErbB family (Lewkowicz, Tchorzewski, Dytnerska, Banasik, & Lewkowicz, 2005), particularly ErbB2 at low levels (Petryszak et al., 2016) and we show that they are phosphorylated and regulated following exposure to inflammatory stimuli. ErbBs have known roles in suppressing apoptosis of epithelial cells and keratinocytes, but this study is the first to show a role for ErbBs in survival signalling of myeloid cells. Little is known about the roles of ErbBs in neutrophil function. Erbstatin has been shown to inhibit neutrophil ROS production (Dreiem, Myhre, & Fonnum, 2003; Mocsai et al., 1997; Reistad, Mariussen, & Fonnum, 2005) and chemotactic responses (Yasui, Yamazaki, Miyabayashi, Tsuno, & Komiyama, 1994). Other kinase families have been found to play a role in neutrophil survival and neutrophilic inflammation, most notably the cyclin-dependent kinases (CDKs) (Rossi et al., 2006). In accordance with this, compounds targeting CDKs were identified as drivers of neutrophil apoptosis in both our primary and secondary screens. Moreover, p38 MAPK inhibitor compounds were also identified in both zebrafish and human screens, and since this kinase is known to mediate survival signals, these findings give confidence to the robustness of the screen design and execution.

The engagement of apoptosis by the ErbB inhibitors erbstatin and tyrphostin AG825 was confirmed both biochemically by phosphatidylserine exposure, and mechanistically by the caspase inhibitor Q-VD-OPh and loss of Mcl-1. This suggests that inhibiting ErbBs as a therapeutic strategy may achieve an overall anti-inflammatory effect in *in vivo* systems, facilitating clearance by macrophages. In support of this, we provide evidence of increased efferocytosis *in vivo* following tyrphostin AG825 treatment, with no evidence of secondary neutrophil necrosis due to overwhelming macrophage clearance capacity, evidenced both morphologically and by TO-PRO-3 staining.

The ability of ErbB inhibitors to promote neutrophil apoptosis even in the presence of multiple pro-survival stimuli emphasises the potential of ErbB inhibitors in the lung, at sites where inflammatory mediators are in abundance and where neutrophils are exposed to microorganisms. This is supported by the ability of tyrphostin AG825 to prevent early pro-survival signalling in response to GMCSF, including the phosphorylation of AKT. This precedes the onset of apoptosis, occurring at a time point (15 min) where apoptosis is typically less than 1%. Others have shown the ability of erbstatin to prevent GMCSF-mediated activation of PI3K in human neutrophils, although the impact on cell survival was not studied (al-Shami, Bourgoin, & Naccache, 1997). Therefore, ErbBs may function as an early and upstream component of the survival pathway in neutrophils. Subsequent impact on Mcl-1 destabilisation by tyrphostin AG825 at 8h suggests a cellular mechanism by which these pro-apoptotic effects are mediated.

The effects of ErbB inhibitors in driving spontaneous apoptosis suggest that, under certain circumstances, ErbB activity might be required for constitutive neutrophil survival. It is not clear what, if anything, engages ErbB signalling in culture. The rapid phosphorylation of ErbB2 following N^6^-MB-cAMP treatment (30 min) suggests that perhaps a ligand is not required, or that the neutrophils can rapidly release ErbB agonists in an autocrine manner. Unlike all other ErbBs, ErbB2 monomers exist in a constitutively active conformation and can form homodimers that do not require a ligand for activation (Fan et al., 2008). ErbBs achieve great signalling diversity: in part because of the individual biochemical properties of ligands and multiple homo-heterodimer combinations, and in part because they activate multiple components including those known to be critical in neutrophil cell survival such as PI3K, MAPK and GSK-3, as well as phosphorylating the Bcl-2 protein Bad which inhibits its death-promoting activity (Yarden & Sliwkowski, 2001).

A limitation of our study is the genetically intractability of human neutrophils, meaning we cannot exclude the possibility that the inhibitors are having off target effects in this system. Mammalian models of ErbB deletion are limited by profound abnormalities in utero and during development (Britsch et al., 1998; Dackor, Strunk, Wehmeyer, & Threadgill, 2007; Gassmann et al., 1995; Miettinen et al., 1995; Riethmacher et al., 1997). For this reason, CRISPR/Cas9 was used to knockdown *egfra* and *erbb2* in zebrafish, which confirmed a role for ErbBs in resolving inflammation. Targeting ErbBs genetically and pharmacologically reduces the number of neutrophils at the site of injury in zebrafish, which may reflect inhibition of a number of pathways that regulate neutrophil number in the tissue, including migration pathways (Ellett, Elks, Robertson, Ogryzko, & Renshaw, 2015). However, the increase in apoptotic neutrophil count at the site of injury with ErbB inhibitor treatment suggests ErbBs may be inducing anti-apoptotic signalling pathways within this inflammatory environment, which could at least in part be causing the phenotype. The reduced neutrophil count at the injury site may also be due to the increase in apoptotic neutrophils in the CHT, which may be preventing neutrophil migration to sites of injury. The unchanged whole body neutrophil number is potentially due to compensatory upregulation of neutrophil production within the CHT. Genetic deletion, but not pharmacological inhibition, of *egfra* and *erbb2* significantly reduced whole body neutrophil number, which may reflect crispants being without *egfra* and *erbb2* genes from a one-cell stage. Reduced neutrophils at the injury site of crispants may be explained by their reduced whole body neutrophil number, but potentially also defects in the migratory response of these neutrophils to a site of inflammation. Murine models of inflammatory disease, where tyrphostin AG825 was administered either at the time of inflammatory stimulus or once inflammation was established, show an impact on cell number, proinflammatory cytokine production and neutrophil apoptosis, further validating the use of ErbB inhibitors to reduce inflammation.

In conclusion, we have identified a previously undefined role for ErbB RTKs in neutrophil survival pathways and a potential new use for ErbB inhibitors in accelerating inflammation resolution. These findings suggest the ErbB family of kinases may be novel targets for treatments of chronic inflammatory disease, and the potential for repurposing ErbB inhibitors currently in use for cancer may have significant clinical potential in a broader range of indications.

## Materials and Methods

### Experimental design

Our objectives for this study are to identify compounds that are able to resolve neutrophilic inflammation. To do this we performed unbiased chemical screens in both human neutrophils *in vitro* and zebrafish models of inflammation *in vivo.* Results were validated in murine models of peritoneal and airway inflammation and zebrafish tail injury models. Genetic evidence was obtained by CRISPR/Cas9 genetic editing in zebrafish.

### Isolation and culture of human neutrophils

Neutrophils were isolated from peripheral blood of healthy subjects and COPD patients by dextran sedimentation and discontinuous plasma-Percoll gradient centrifugation, as previously described (Haslett, Guthrie, Kopaniak, Johnston, & Henson, 1985; I. Ward, Dransfield, Chilvers, Haslett, & Rossi, 1999) in compliance with the guidelines of the South Sheffield Research Ethics Committee (for young healthy subjects; reference number: STH13927) and the National Research Ethics Service (NRES) Committee Yorkshire and the Humber (for COPD and age-matched healthy subjects; reference number: 10/H1016/25).

Informed consent was obtained after the nature and possible consequences of the study were explained. Mean age in years was 61.7±2.3 (n=10) and 66.0±3.6 (n=7) for COPD and age-matched healthy subjects respectively. Ultrapure neutrophils, for Kinexus antibody array experiments, were obtained by immunomagnetic negative selection as previously described (Sabroe, Jones, Usher, Whyte, & Dower, 2002). Neutrophils were cultured (2.5x10^6^/ml) in RPMI 1640 (Gibco, Invitrogen Ltd) supplemented with 10% FCS 1% penicillin-streptomycin, in the presence or absence of the following reagents: GMCSF (PeproTech, Inc), N^6^-MB-cAMP (Biolog), anti-ErbB3 blocking antibody, Tyrphostin AG825 (both Sigma-Aldrich), CP-724714 (AdooQ Bioscience), Erbstatin analog (Cayman Chemicals), Pyocyanin (Usher et al., 2002) or compounds from PKIS (Published Kinase Inhibitor Set 1, GlaxoSmithKline) at concentrations as indicated.

*In vitro screening of PKIS in neutrophil apoptosis assays*. PKIS consists of 367 small molecule protein kinase inhibitors and is profiled with respect to target specificity (Elkins et al., 2016). In primary screen experiments, neutrophils (from 5 independent donors over 5 days) were incubated with each compound at 62µM for 6h. Apoptosis was measured by flow cytometry (Attune, Invitrogen). Secondary screening was performed with selected compounds that accelerated neutrophil apoptosis greater than twofold in the primary screen. Compounds were incubated with neutrophils at 10µM for 6h and apoptosis assessed by Attune flow cytometry.

### Human neutrophil apoptosis assays

Neutrophil apoptosis was assessed by light microscopy and by flow cytometry. Briefly, for the assessment of apoptosis by light microscopy based on well-characterised morphological changes, neutrophils were cytocentrifuged, fixed with methanol, stained with Reastain Quick-Diff (Gentaur), and then apoptotic and non-apoptotic neutrophils were counted with an inverted, oil immersion microscope (Nikon Eclipse TE300, Japan) at 100X magnification (Savill et al., 1989). To assess apoptosis by flow cytometry, neutrophils were stained with PE conjugated Annexin-V (BD Pharminogen) and TO-PRO-3 (Thermofisher Scientific) (Savill et al., 1989; Vermes, Haanen, Steffens-Nakken, & Reutelingsperger, 1995; C. Ward et al., 1999) and sample acquisition was performed by an Attune flow cytometer (Life Technologies) and data analysed by FlowJo (FlowJo LLC).

### Kinexus antibody array

Neutrophils were incubated with N^6^-MB-cAMP [100µM] for 30 and 60 min or lysed immediately following isolation (t0). Cells were lysed in PBS containing Triton-X, 1µM PMSF and protease inhibitor cocktail and following 2 min on ice were centrifuged at 10,000 RPM to remove insoluble material. Lysates (containing protein at 6mg/mL) from four donors were pooled prior to Kinex antibody microarray analysis (Kinexus Bioinformatics) (H. Zhang & Pelech, 2012). Lysates are subjected to 812 antibodies including phospho-site specific antibodies to specifically measure phosphorylation of the target protein. Fluorescent signals from the array were corrected to background and log2 transformed and a Z score calculated by subtracting the overall average intensity of all spots within a sample, from the raw intensity for each spot, and dividing it by the standard deviations (SD) of all the measured intensities within each sample (Cheadle, Vawter, Freed, & Becker, 2003). Z ratio values are further calculated by taking the difference between the averages of the Z scores and dividing by the SD of all differences of the comparison (e.g, 30 min treated samples versus 0 min control). A Z ratio of ±1.5 is considered to be a significant change from control.

### Western blotting

Whole cell lysates were prepared by resuspending human neutrophils (5X10^6^) in 50µl hypotonic lysis buffer (1mM PMSF, 50mM NaF, 10mM Sodium orthovanadate, protease inhibitors cocktail in water), and by boiling with 50µl 2X SDS buffer (0.1M 1,4-Dithio-DL-threitol, 4% SDS, 20% Glycerol, 0.0625M Tris-HCl pH6.8 and 0.004% Bromophenol blue). Protein samples were separated by SDS-polyacrylamide gel electrophoresis, and electrotransfer onto PVDF (polyvenylidene difluoride) membranes was performed by semi-dry blotting method. Membranes were then blocked with 5% skimmed milk in TBS-tween and probed against antibodies to p-AKT (Cell Signalling Technology), AKT (Cell signalling Technology), Mcl-1 (Santa Cruz Biotechnology) or p38 (loading control, StressMarq Biosciences Inc.), followed by HRP-conjugated secondary antibodies and detection with chemiluminescent substrate solution ECL2 (GE Healthcare).

### Fish husbandry

The neutrophil-specific, GFP-expressing transgenic zebrafish line, *Tg(mpx:GFP)i114*, (referred to as *mpx:*GFP) (Renshaw et al., 2006) was raised and maintained according to standard protocols (Nüsslein-Volhard & Dahm, 2002) in UK Home Office approved aquaria in the Bateson Centre at the University of Sheffield, according to institutional guidelines. Adult fish are maintained in 14h light and 10h dark cycle at 28°C.

### Zebrafish tail injury model of inflammation

*PKIS screening*: Tail fin transection was performed on *mpx:*GFP zebrafish larvae at 3 days post-fertilisation (dpf) (Elks, Loynes, & Renshaw, 2011; Renshaw et al., 2006). At 6h post-injury (hpi), larvae that had mounted a good inflammatory response, as defined by recruitment of >15 neutrophils to the injury site, were arrayed at a density of 3 larvae per well and incubated with PKIS compounds at a final concentration of 25µM or vehicle control for a further 6h. At 12 hpi, the plate was scanned using prototype PhenoSight equipment (Ash Biotech). Images were scored manually as described previously (Robertson et al., 2014). In brief, each well of three larvae was assigned a score between 0-3, corresponding to the number of larvae within the well with a reduced number of neutrophils at the site of injury. Kinase inhibitors which reduced green fluorescence at the injury site to an extent that their mean score was ≥1.5 were regarded as hit compounds.

*ErbB inhibition studies:* Briefly, 2 dpf *mpx:*GFP larvae were treated with Tyrphostin AG825 [10µM] for 16h before undergoing tailfin transection (Elks et al., 2011; Renshaw et al., 2006). The number of neutrophils at the site of injury was determined at 4 and 8 hpi by counting GFP-positive neutrophils by fluorescent microscopy. To enumerate neutrophils across the whole body, uninjured larvae were treated with Tyrphostin AG825 [10µM] for 24h and then mounted in 0.8% low-melting point agarose (Sigma-Aldrich) followed by imaging by fluorescence microscopy (Nikon Eclipse TE2000-U) at 4X magnification, followed by manual counting.

### Zebrafish Apoptosis Assays

Larvae from each experimental group were pooled into 1.5mL eppendorf tubes. TSA signal amplification of GFP-labelled neutrophils (driven by endogenous peroxidase activity) was carried out using TSA® Plus Fluorescein System (Perkin Elmer). Larvae were fixed overnight in 4% paraformaldehyde at 4°C after which they were subjected to proteinase K digestion. Larvae were post-fixed in 4% paraformaldehyde, before subsequent TUNEL staining for apoptosis using ApopTag® Red In Situ Apoptosis Detection Kit (Millipore).

Larvae were then mounted in low-melting point agarose and images acquired and analysed using UltraVIEWVoX spinning disc confocal laser imaging system with Volocity® 6.3 software (Perkin Elmer). Apoptotic neutrophil count was determined firstly by identifying cells with co-localisation of the TSA and TUNEL stains, then confirmed by accounting for apoptotic neutrophil morphology.

### Generation of transient CRISPR/Cas9 zebrafish mutants

Transient dual knockdown of *egfra* and *erbb2* was induced using a Cas9 nuclease (New England Biolabs) in combination with transactivating RNA (tracr) and synthetic guide RNAs specific to zebrafish *egfra* and *erbb2* genes (Merck). Tyrosinase guide RNA was used as a control as described previously (Isles et al., 2019). Guide RNAs were designed using the online tool CHOPCHOP (https://chopchop.cbu.uib.no/) with the following sequences: *efgra*: TGAATCTCGGAGCGCGCAGGAGG; *erbb2*: AACGCTTTGGACCTACACGTGGG; *tyrosinase*: GGACUGGAGGACUUCUGGGG. Each guide RNA was resuspended to 20μM in nuclease-free water with 10mM Tris-HCl (pH8). Guide RNA [20μM], tracr [20μM] and Cas9 protein [20μM] were combined (in a 1:1:1 ratio). 0.5μL phenol red was added to each injection solution for visualisation. A graticule was used to calibrate glass capillary needles to dispense 0.5nL of injection solution, and 1nL was injected into the yolk sac of single-cell stage *mpx*:GFP embryos. Tail injury assays were carried out at 2 dpf as described above.

### Genotyping of crispant larvae

High-resolution melt curve analysis was used to determine the rate of *egfr* and *erbb2* mutation in larvae at 2 dpf. Genomic DNA was collected from individual larvae in both the control and experimental groups, by adding 90 μL 50 mM NaOH to each larvae in a 96-well qPCR plate and incubating at 95°C for 20 minutes. 10 μL Tris-HCl (pH 8) was then added as a buffer. Master mixes containing either *egfra* or *erbb2* primers (Integrated DNA Technologies) (sequences in table below) were made up, with each well containing: 0.5 μL 10 μM forward primer, 0.5 μL 10 μM reverse primer, 5 μL 2X DyNAmo Flash SYBR Green (Thermo Scientific), 3 μL milliQ water. 1 μL genomic DNA was added to each master mix in a 96-well qPCR plate. Melt curve analysis was performed and analysed with Bio-Rad Precision Melt Analysis software. Mutation rate was calculated by determining the percentage of *egfra erbb2* larvae that showed a different melt-curve profile to the genomic DNA collected from *tyrosinase* fish (based on 95% confidence intervals).

Primer sequences used for high-resolution melt curve analysis.

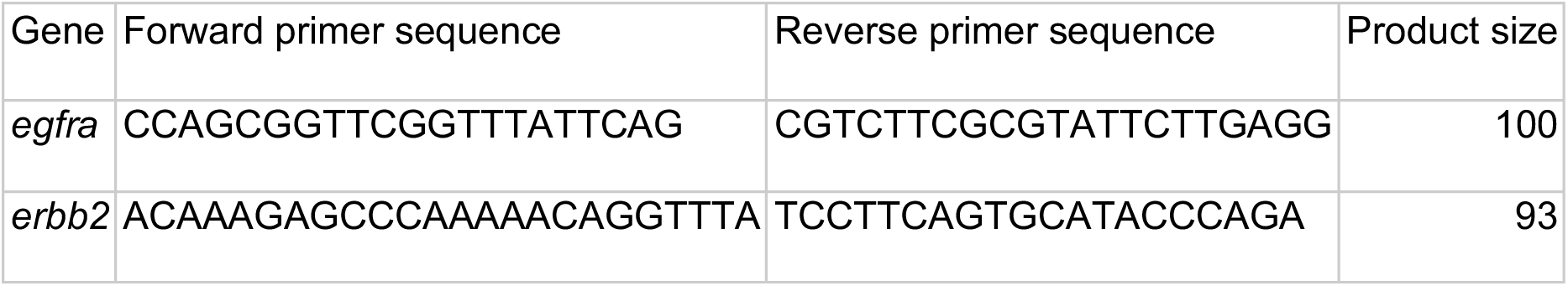

### Murine model of LPS induced acute lung inflammation

C57BL/6 mice (female, 9-10 weeks old) were nebulised with LPS (3 mg per group of 8 mice) (*Pseudomonas aeruginosa*, Sigma-Aldrich) and immediately injected intraperitoneally (i.p.) with either Tyrphostin AG825 (Tocris Bioscience) at 20mg/Kg in 10% DMSO v/v in vegetable oil (8 mice, treatment group) or an equivalent volume of 10% DMSO v/v in vegetable oil (8 mice, control group) (Kedrin et al., 2009; Roos, Berg, Ahlgren, Grunewald, & Nord, 2014). After 48h the mice were sacrificed by terminal anaesthesia by i.p. pentobarbitone and subjected to bronchoalveolar lavage (BAL, 4 x 1mL of saline). BAL samples were microcentrifuged and the cellular fraction counted by a hemocytometer and cytocentrifuged. Neutrophil apoptosis and macrophage efferocytosis of apoptotic neutrophils was quantified by oil immersion light microscopy (Nikon Eclipse TE300, Japan).

### Murine model of zymosan-induced peritonitis

C57BL/6 mice were i.p. injected with 1mg zymosan (Sigma-Aldrich) and 4h later injected i.p with 20mg/Kg Tyrphostin AG825 in 10% DMSO v/v in vegetable oil (5 mice, treatment group) or an equivalent volume of 10% DMSO v/v in vegetable oil (5 mice, control group). At 20h the mice were subjected to terminal gaseous anaesthesia (isoflurane) followed by a cardiac puncture and peritoneal lavage (4 x 1mL of saline). WBC, neutrophils and macrophages were enumerated in blood by an automated haematology analyser (KX-21N, Sysmex, Milton Keynes, UK). Lavage samples were microcentrifuged and the cellular fraction subjected to flow cytometry and cytocentrifuged for light microscopy. IL-6, KC (Duoset ELISA kits, R&D systems) and IgM (Thermofisher Scientific) in cell free lavage were measured by ELISA as per manufacturer’s instructions.

### Statistical analysis

Data were analysed using GraphPad Prism 8 (GraphPad Software, San Diego, CA) using one-way or two-way ANOVA (with appropriate post-test) for all *in vitro* data and appropriate *in vivo* experiments. Non-parametric tests (Mann-Whitney U-test or Kruskal-Wallis test) were used for selected *in vivo* experiments with non-Gaussian distribution. Data are expressed as mean ± SEM (standard error of mean), and significance was accepted at p<0.05.

## Acknowledgements

**General:** We thank Lynne Williams, Carl Wright, Jessica Willis, Elizabeth Marsh and Catherine Loynes for help with animal experiments as well as volunteers and patients who donated blood to this study. We thank the Bateson Centre aquaria staff for their assistance with zebrafish husbandry.

## Author contributions

AR, KMH, IS, DHD, MKBW, SAR and LRP designed experiments and analysed data. AR, KMH, KDH, HMI, AER, NK, DS, C Tabor, C Tulotta, AART and KH performed experiments. AR, SAR, KMH and LRP wrote the manuscript. All authors contributed intellectual input to the concept of the study and to the writing and editing of this manuscript.

## Supplementary Materials

**Fig. S1.**
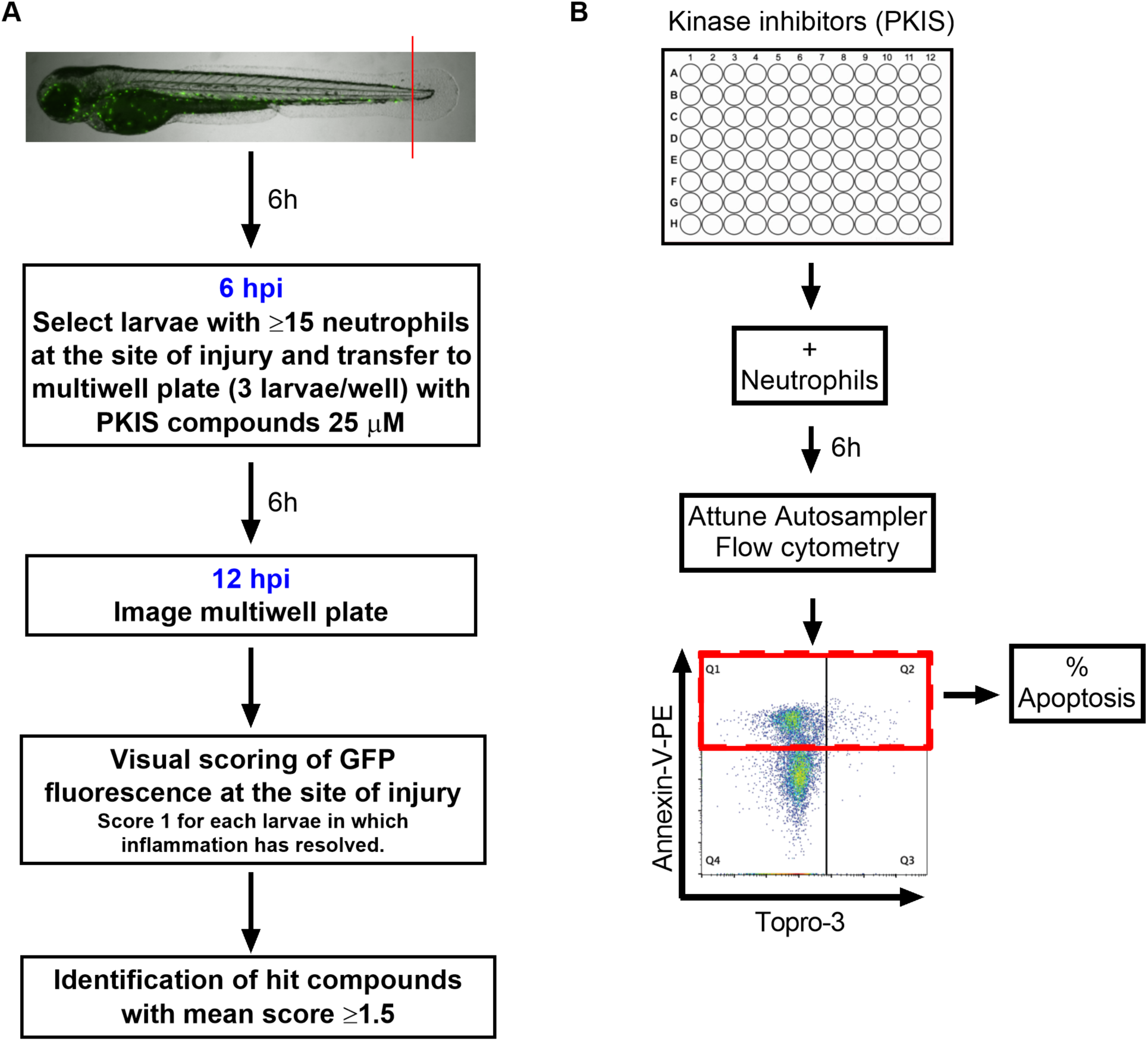
Schematics showing PKIS screen design. (A) Tail fin transected 3 dpf Tg(mpx:GFP)i114 zebrafish larvae that had generated an inflammatory response at 6 hpi were incubated with individual PKIS compounds [25µM] for a further 6h. Larvae were imaged and manually scored between 0-3 on the basis of green fluorescence at the injury site. (B) PKIS compounds were incubated with primary human neutrophils for 6h. Apoptosis was assessed by Annexin V/TO-PRO-3 staining by flow cytometry and the percentage apoptosis calculated as Annexin V single plus Annexin V/TO-PRO-3 dual positive events (as indicated by red box).

**Fig. S2.**
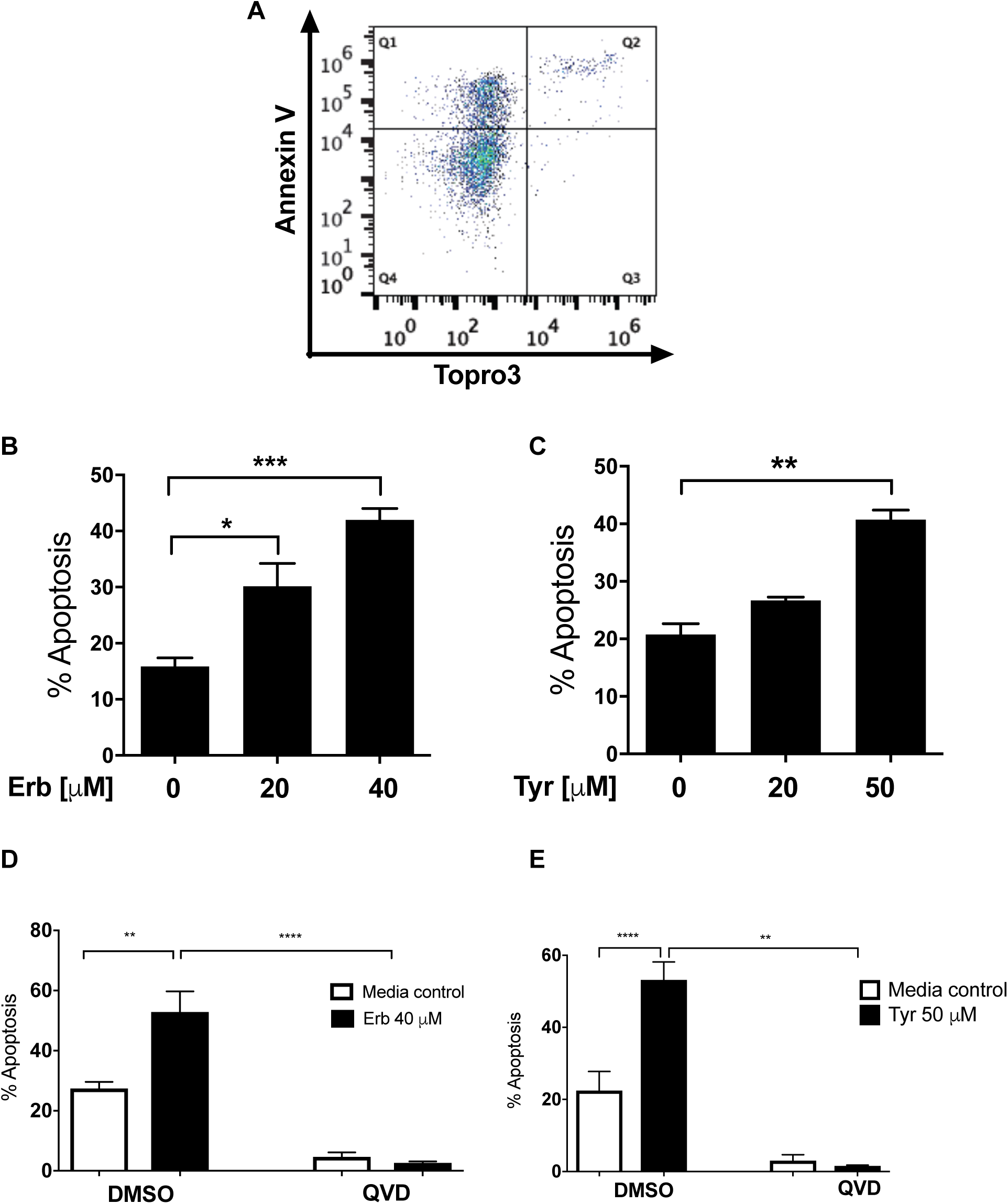
Erbstatin and tyrphostin AG825 induces caspase-dependent neutrophil apoptosis. Neutrophils were incubated with DMSO or 20µM or 40µM erbstatin (Erb, A&B) or tyrphostin AG825 (Tyr, C) for 6h. Apoptosis was assessed by Annexin V/TO-PRO-3 staining by flow cytometry and the percentage apoptosis calculated as Annexin V single plus Annexin V/TO-PRO-3 dual positive events. (A) Representative quadrant plot of Erbstatin-treated neutrophils showing distribution of Annexin V and TO-PRO-3 positive events. (D-E) Neutrophils were incubated with DMSO or Erbstatin [40µM] in the presence or absence of the pan caspase inhibitor, Q-VD-OPh [1µM] for 6h (D) or 20h (E). Apoptosis was assessed by light microscopy. The data are expressed as mean percentage apoptosis ± SEM from 3 (B), 4 (C&E), or 5 (D) independent experiments. Statistical differences were calculated by ANOVA (with Dunnett’s (B-C) Bonferroni’s (D-E) post-tests) and indicated as *p≤0.05, **p≤0.01, ***p≤0.001, ****p≤0.0001.

**Fig. S3.**
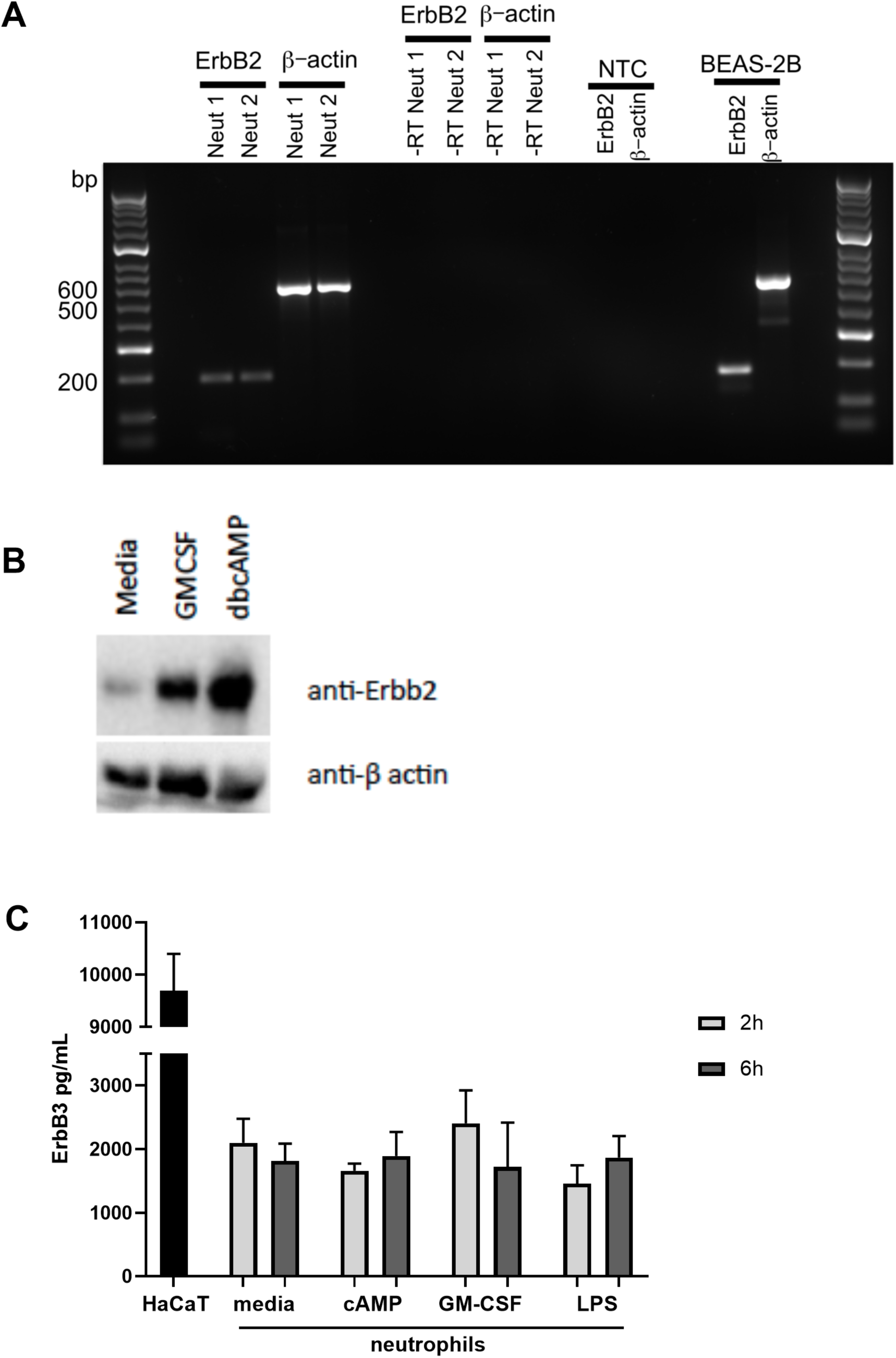
ErbB2 and ErbB3 expression and regulation in human neutrophils. ErbB2 was detected in neutrophils and the positive control cell line, BEAS-2B, by RT-PCR (A). Neutrophils were treated with GMCSF [50u/mL] and dbcAMP [10μM] for 5 hours and lysates subjected to SDS PAGE. Membranes were immunoblotted with antibodies to ErbB2 antibody or β-actin as a loading control. A 60kD band was detected (lower molecular weight ErbB family products are well-documented (Jackson, Browell, Gautrey, & Tyson-Capper, 2013), which was upregulated by GMCSF and dbcAMP. NTC – no template control. The image is representative of three independent experiments (B). ErbB3 was detected by ELISA in human neutrophils and the positive control cell line, HaCaT. Neutrophils were treated with media, dbcAMP [500µM], GM-CSF [50u/mL] or LPS [1µg/mL] for 2h or 6h, after which lysates were collected and ELISA detecting total human ErbB3 was carried out. N=4 healthy human neutrophil donors. Bars indicate mean + SEM. (C).

**Table S1.**
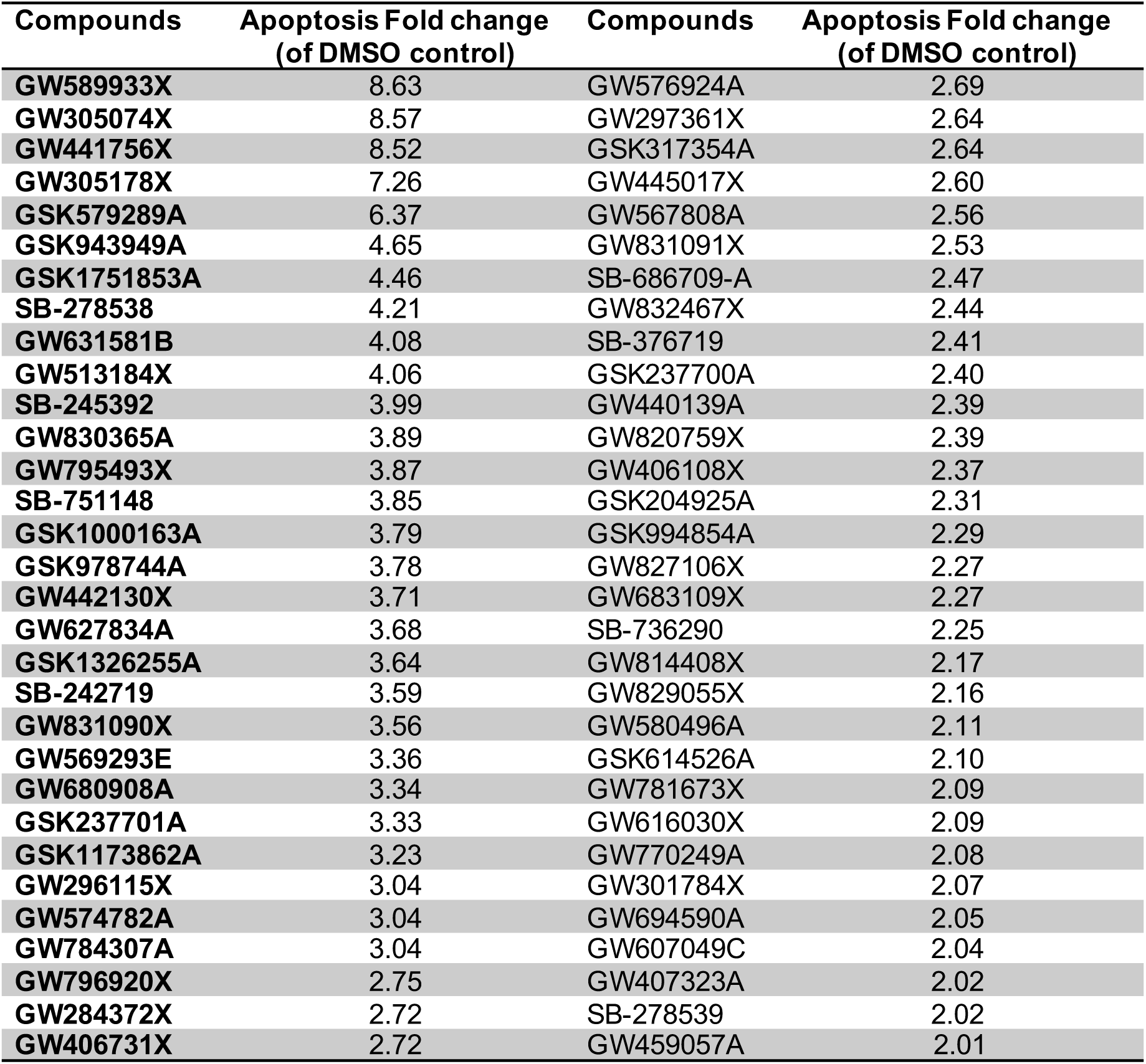
PKIS compounds that accelerated neutrophil apoptosis >2 fold over control. PKIS compounds were incubated with neutrophils for 6h and apoptosis was assessed by Annexin V/TO-PRO-3 staining by flow cytometry. Sixty-two compounds accelerated apoptosis ≥ 2 fold and compound names are presented here, along with fold change over control.

| Compounds | Apoptosis Fold change<br>(of DMSO control) | Compounds | Apoptosis Fold change<br>(of DMSO control) |
| --- | --- | --- | --- |
| GW589933X | 8.63 | GW576924A | 2.69 |
| GW305074X | 8.57 | GW297361X | 2.64 |
| GW441756X | 8.52 | GSK317354A | 2.64 |
| GW305178X | 7.26 | GW445017X | 2.60 |
| GSK579289A | 6.37 | GW567808A | 2.56 |
| GSK943949A | 4.65 | GW831091X | 2.53 |
| GSK1751853A | 4.46 | SB-686709-A | 2.47 |
| SB-278538 | 4.21 | GW832467X | 2.44 |
| GW631581B | 4.08 | SB-376719 | 2.41 |
| GW513184X | 4.06 | GSK237700A | 2.40 |
| SB-245392 | 3.99 | GW440139A | 2.39 |
| GW830365A | 3.89 | GW820759X | 2.39 |
| GW795493X | 3.87 | GW406108X | 2.37 |
| SB-751148 | 3.85 | GSK204925A | 2.31 |
| GSK1000163A | 3.79 | GSK994854A | 2.29 |
| GSK978744A | 3.78 | GW827106X | 2.27 |
| GW442130X | 3.71 | GW683109X | 2.27 |
| GW627834A | 3.68 | SB-736290 | 2.25 |
| GSK1326255A | 3.64 | GW814408X | 2.17 |
| SB-242719 | 3.59 | GW829055X | 2.16 |
| GW831090X | 3.56 | GW580496A | 2.11 |
| GW569293E | 3.36 | GSK614526A | 2.10 |
| GW680908A | 3.34 | GW781673X | 2.09 |
| GSK237701A | 3.33 | GW616030X | 2.09 |
| GSK1173862A | 3.23 | GW770249A | 2.08 |
| GW296115X | 3.04 | GW301784X | 2.07 |
| GW574782A | 3.04 | GW694590A | 2.05 |
| GW784307A | 3.04 | GW607049C | 2.04 |
| GW796920X | 2.75 | GW407323A | 2.02 |
| GW284372X | 2.72 | SB-278539 | 2.02 |
| GW406731X | 2.72 | GW459057A | 2.01 |

